# Sexing carnivores from faecal metagenomes: calibrating *R*_*x*_ for stone marten and red fox

**DOI:** 10.64898/2026.11.04.764345

**Authors:** Vesna Vujić, Csilla Mayer, Georgina Helyes, Ákos Maróti-Agóts, Gergely Maróti, István Csabai, Norbert Solymosi

**Affiliations:** Centre for Bioinformatics, University of Veterinary Medicine, Budapest, 1078, Hungary; Institute of Animal Breeding Nutrition and Laboratory Animal Science, University of Veterinary Medicine, Budapest, 1078, Hungary; Institute of Plant Biology, HUN-REN Biological Research Center, Szeged, 6726, Hungary; Department of Physics of Complex Systems, Eötvös Loránd University, Budapest, 1117, Hungary

**Keywords:** molecular sexing, *R*_*x*_, *R*_*y*_, low-coverage sequencing, non-invasive sampling, competitive mapping, human contamination, Y chromosome assembly, *Martes foina*, *Vulpes vulpes*

## Abstract

Knowing the sex of the animal that deposited a scat adds value to non-invasive monitoring, but read-ratio sexing methods were developed for human ancient DNA. We examined their use in shotgun metagenomes of seven stone marten (*Martes foina*) and red fox (*Vulpes vulpes*) scats. Human DNA, present in every library, was removed by competitive mapping against the host assembly and the human reference. Applying the Skoglund *R*_*y*_ metric to the marten samples yielded 0 to 25 Y-mapping reads per sample and failed to identify either of the two animals that *R*_*x*_ places firmly in the male range, so we adopted the X-to-autosome ratio *R*_*x*_ of Mittnik et al. instead, reimplemented as an open-source, species-agnostic Python tool. To test whether its human-derived thresholds transfer, we simulated female and male genomes of both species and drew 200 replicate subsamples at each of fourteen depths from 1,000 to 100,000 mapped reads. Nine of 11,200 replicates were assigned to the wrong sex, all of them below 2,000 mapped reads; above that depth the only failure was non-assignment, in under 5% of replicates above 7,500 reads and under 1% above 15,000. Applied to the scats, *R*_*x*_ identified four females and two males; one sample with 596 host-assigned reads was left unassigned. With competitive removal of human reads, *R*_*x*_ and its original thresholds are usable in these carnivores.

## 1 Introduction

The sex of the animal that deposited a scat is one of the few pieces of individual-level information that non-invasive sampling could deliver cheaply, and sex ratios and sex-specific dispersal are central to any demographic interpretation of scat-based survey data [1–4]. Conventional molecular sexing amplifies sex-linked markers [5], which requires species-specific primers and performs poorly on the short, degraded templates typical of field-collected faeces [6–8]. Shotgun sequencing needs no primers: because females carry two X chromosomes and males one, the proportion of reads mapping to the X relative to the autosomes differs systematically between the sexes, and the same logic applied to the Y gives a complementary statistic.

Both statistics in general use were developed for human ancient DNA. *R*_*y*_ [9] is the ratio of Y-mapping reads to the sum of X- and Y-mapping reads; *R*_*x*_ [10] compares X-mapping reads to the reads on each autosome after normalising for chromosome length. Their thresholds were calibrated on human data against human assemblies, and neither has an established operating range in non-model mammals. The metrics have nonetheless been transferred to other taxa, for example to ancient elephant remains [11], each transfer carrying the implicit assumption that thresholds derived from one karyotype and one assembly apply to another. Dedicated pipelines exist for identifying sex chromosomes in non-model species [12], but they are designed for genome-scale data rather than for the few thousand host reads a faecal metagenome typically yields.

Carnivore faecal metagenomes add two complications (Figure 1). The first is human DNA: scats collected along hiking trails are handled by people and lie where people frequent, and because the human and carnivore genomes share large conserved regions – X-linked regions among them – reads of human origin map to a carnivore reference offered alone, biasing the quantity both metrics measure. Competitive mapping against a combined host-plus-human reference is the standard remedy [13–15]; reference bias acts on read counts in the same way [16]. The second is the state of carnivore Y assemblies. The mammalian Y is dominated by ampliconic and heterochromatic sequence that short-read assemblers cannot resolve, and it was the last part of the genome completed even in humans [17–20]. Here the consequence is concrete: the *Martes foina* reference genome mMarFoi2.1 [21] carries an annotated Y of only 5.6 Mb, and VulVul3 of *Vulpes vulpes* [22] none at all, even though the red fox Y has been characterised by other means [23].

**Figure 1.**
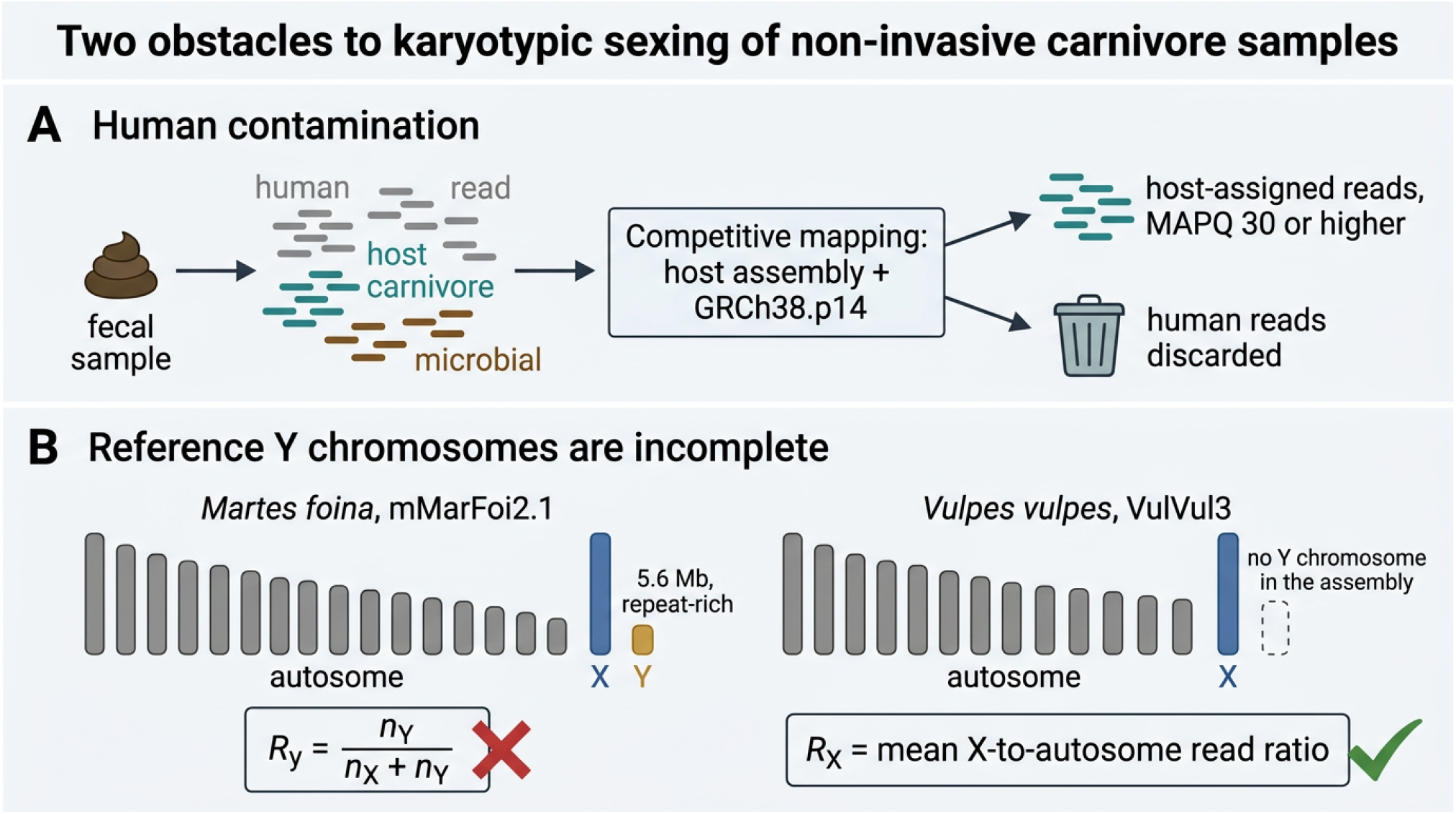
Human contamination and incomplete Y assemblies. (A) Faecal metagenomes contain host, microbial and human DNA; mapping competitively against the host assembly together with GRCh38.p14 assigns human reads away from the host genome, and only host-assigned reads with MAPQ ≥ 30 enter the ratio calculation. (B) The Y chromosome is a weak denominator in these species: the *M. foina* assembly mMarFoi2.1 contains a short, repeat-rich Y and the *V. vulpes* assembly VulVul3 contains none, which disqualifies *R*_*y*_ and favours the X-to-autosome ratio *R*_*x*_. The panel is a schematic illustration and contains no measured values; the measured calibration results are shown in Figure 3.

We applied both metrics to seven non-invasively collected carnivore scats. *R*_*y*_ proved unusable, as the Y assemblies anticipate, so we turned to *R*_*x*_, reimplemented it as a species-agnostic tool that needs no Y chromosome, and calibrated it by simulation for both species before reading the empirical values against the result. The calibration also established the minimum depth at which an assignment is supported, which turns out to matter for one of the seven samples. The samples themselves and the broader biological analyses are described in two companion manuscripts.

## 2 Materials and methods

### 2.1 Samples and sequencing

Seven visibly fresh carnivore scats were collected from stones along hiking trails in two Hungarian mid-mountain ranges in August and September 2025. Total DNA was extracted with the Quick-DNA Fecal/Soil Microbe Kit (Zymo Research) in a dedicated, HEPA-filtered and UVC-equipped laboratory; paired-end libraries were prepared with the NEBNext Ultra II DNA Library Prep Kit and sequenced on an Illumina NextSeq instrument (TG NextSeq 500/550 High Output Kit v2, 300 cycles), yielding 4.0–5.0 million cleaned read pairs per sample after trimming with TrimGalore (v0.6.6) [24]. Full details of collection, extraction and sequencing are given in the companion molecular-ecology manuscript. Mitogenomic analysis assigned five samples to *M. foina* (1, 3, 4, 6, 7) and two to *V. vulpes* (2, 5).

### 2.2 Competitive mapping and read filtering

Because the samples were contaminated with human DNA to varying degrees, sex estimation was based on competitive mapping. For the marten-derived samples the combined reference comprised the *M. foina* assembly mMarFoi2.1 [21] and the human assembly GRCh38.p14 [25, 26]; for the fox-derived samples it comprised the *V. vulpes* assembly VulVul3 [22, 27] and the same human assembly. Alignment used Bowtie2 (v2.4.5) [28] in –very-sensitive-local mode. Only reads mapping to a host chromosome with MAPQ 30 entered the ratio calculations, a threshold that excludes reads placed ambiguously between the two references or within repetitive regions. We took counts read by read from the alignment files rather than from samtools idxstats, which counts every mapped read regardless of mapping quality. Coverage statistics were obtained with SAMtools [29].

### 2.3 The *R*_*y*_ statistic

Before adopting *R*_*x*_ we applied the *R*_*y*_ statistic of Skoglund et al. [9] to the five marten samples, using the reference implementation (https://github.com/pontussk/ry_compute) and the published thresholds (*R*_*y*_ *>* 0.075 male, *R*_*y*_ *<* 0.016 female). The calculation used the same competitively mapped, MAPQ-filtered alignments as the *R*_*x*_ estimates, so that the two statistics differ only in which chromosomes they count. *R*_*y*_ was not applicable to the fox samples, because VulVul3 contains no Y chromosome.

### 2.4 The *R*_*x*_ statistic and its implementation

We followed the approach of Mittnik et al. [10]: for each autosome *i*, the ratio of X-mapping reads to reads mapping to that autosome is normalised by the corresponding chromosome lengths, and *R*_*x*_ is the mean of these per-autosome ratios. Formally, with *m*_*X*_ reads on the X chromosome of length *s*_*X*_ and *m*_*i*_ reads on autosome *i* of length *s*_*i*_ (*i* = 1, …, *k*),

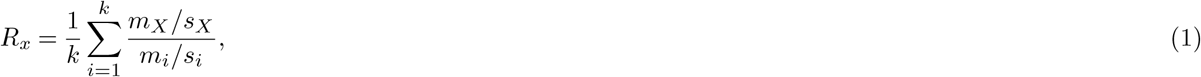

and the 95% confidence interval is *R*_*x*_ *±* 1.96 SE, where SE is the standard error of the mean of the *k* per-autosome ratios. Only the assembled autosomes and the X chromosome enter the calculation. The mitochondrial genome is excluded because, present in many copies per cell, it would act as a strongly inflated reference term and depress *R*_*x*_; unplaced scaffolds are excluded as well.

We reimplemented the published R code [10] as a function karyoratio (v2.0.0), a command-line Python tool, that accepts any assembly with an identified X chromosome, so that it is not tied to the human chromosome set. Another change to Mittnik et al. [10] approach: the correlation between chromosome length and the number of reads mapping to each chromosome, which the original implementation quantifies by linear regression, is quantified here by the Spearman rank correlation coefficient *ρ*. This makes the diagnostic insensitive to the occasional chromosome that attracts a disproportionate number of reads. The coefficient is reported alongside each estimate as a quality indicator: a value close to 1 indicates that read counts scale with chromosome length as expected under even coverage.

Mittnik et al. [10] assigned a sample as male if the upper bound of the 95% CI lies below 0.60 and as female if the lower bound exceeds 0.80. Because these thresholds were established on human samples, we could not assume that they transfer to the species studied here, and we therefore calibrated the metric by simulation. The codes required for this simulation are also available in the karyoratio repository.

### 2.5 Simulation for *R*_*x*_ statistic threshold estimation

For each species, synthetic female and male genomes were constructed from the reference assembly. The synthetic female marten genome contained two copies of every autosome and two copies of the X chromosome; the synthetic male marten genome was identical except that one X copy was replaced by the Y chromosome. Because the fox assembly contains no Y chromosome, the synthetic male fox genome contained two autosome sets and a single X chromosome, with no Y, while the synthetic female fox genome was built as described for the marten.

Each female X is a duplicate of the same haplotype, so the simulation models copy number rather than genetic variability.

Reads were generated with ART [30] under an error profile estimated from our own data by sample 1 for the marten and sample 2 for the fox. Paired 151 bp reads were drawn from each of the four synthetic genomes (two species *×*two sexes) with the fragment-length distribution measured in the same sample (333 *±* 80 bp for the marten, 306 *±* 73 bp for the fox), to a pool of 20,000,000 reads per genome. Each pool was mapped to the corresponding reference assembly alone, without the human genome: every simulated read is host-derived, so the human sequences of a combined reference would remain empty. From the mapped pool, 200 replicate subsamples were drawn at each of fourteen depths – 1,000, 1,500, 1,700, 2,000, 2,200, 2,500, 3,000, 5,000, 7,500, 10,000, 15,000, 25,000, 50,000 and 100,000 mapped reads – and *R*_*x*_ was computed for each by the same implementation as for the empirical samples. Subsampling draws read pairs rather than individual reads, because the two mates of a pair almost always map to the same chromosome and are therefore not independent observations. Depths are reported in mapped reads passing the MAPQ filter, the unit karyoratio reports and the unit of *n* in Table 2; the deepest level is a twentieth of the pool, so replicates within a level overlap little. The design is summarised in Figure 2.

**Figure 2.**
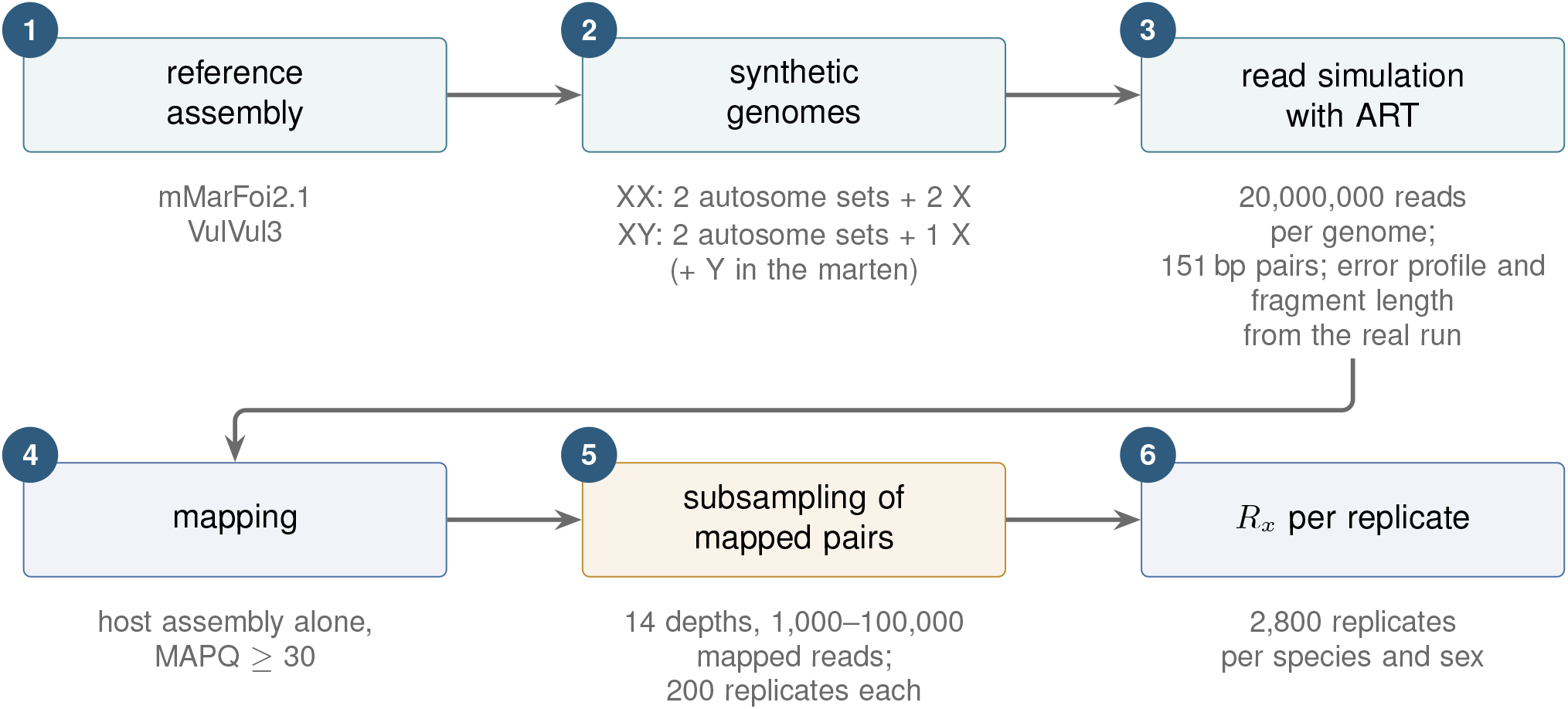
Design of the calibration simulations. Synthetic female and male genomes were assembled from each reference assembly. The reads were generated with ART under the error profile estimated from the real sequencing run. We subsampled the mapped pool at fourteen depths in 200 replicates, and each subsample underwent the same *R*_*x*_ calculation as the empirical samples. The fox “male” genome contains no Y chromosome because none is present in the VulVul3 assembly.

## 3 Results

### 3.1 *R*_*y*_ is uninformative on the marten samples and unavailable for the fox

*R*_*y*_ could not be computed for the two fox samples because the VulVul3 assembly contains no Y chromosome. For martens, Table 1 reports the *R*_*y*_ estimates. Samples 1 and 3 fall at or below the female threshold and sample 4 well below it, sample 6 falls between the thresholds, samples 1 and 7 give intervals that exclude the male range without reaching the female one. The number of Y-mapping reads was 0 to 25 per sample. At these counts, the normal approximation behind the interval breaks down: sample 3 yields *R*_*y*_ = 0 with a standard error of 0, and sample 7 yields an interval with a negative lower bound. No sample reached the male threshold.

**Table 1.** Sex estimates for the *Martes foina* samples obtained with the *R*_*y*_ statistic of Skoglund et al. [9], computed from the same competitively mapped, MAPQ-filtered alignments as the *R*_*x*_ estimates of Table 2. Thresholds: male if *R*_*y*_ *>* 0.075, female if *R*_*y*_ *<* 0.016. nSeqs is the number of reads mapping to the host assembly, and nX and nY the numbers mapping to the X and Y chromosomes; nSeqs exceeds the *n* of Table 2 because it includes the mitochondrial genome and the unplaced scaffolds, which the *R*_*x*_ calculation excludes. No equivalent statistic could be computed for the fox samples, as VulVul3 contains no Y chromosome.

| Sample | nSeqs | nY+nX | nY | *R<sub>y</sub>* | SE | 95% CI | Assignment |
|---|---|---|---|---|---|---|---|
| 1 | 13,919 | 481 | 7 | 0.015 | 0.0055 | 0.004-0.025 | consistent with XX but not XY |
| 3 | 1,220 | 30 | 0 | 0.000 | 0.0000 | 0.000-0.000 | consistent with XX |
| 4 | 350,796 | 17,592 | 25 | 0.001 | 0.0003 | 0.001-0.002 | XX |
| 6 | 15,321 | 416 | 18 | 0.043 | 0.0100 | 0.024-0.063 | Not Assigned |
| 7 | 5,248 | 90 | 2 | 0.022 | 0.0155 | −0.008-0.053 | consistent with XX but not XY |

### 3.2 Simulation-based calibration of *R*_*x*_

Figure 3 summarises the *R*_*x*_ values obtained across the 200 replicates at each simulated depth, separately for each species and sex. The expected separation is present throughout, and the spread of replicate values narrows steadily as depth increases.

**Figure 3.**
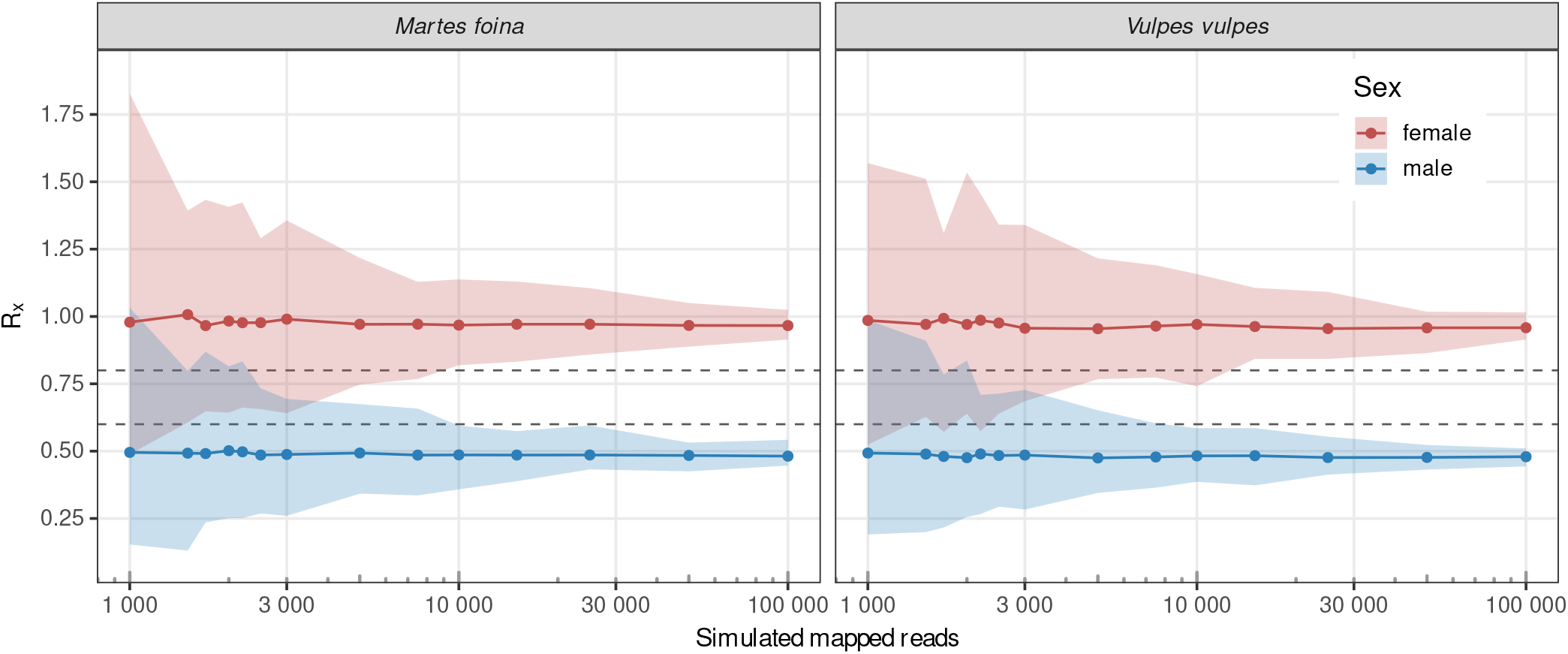
Median (points, joined by a line), minimum and maximum (ranges) of the *R*_*x*_ values calculated for the simulated depths, by species and sex. Each point summarises 200 replicate subsamples. The band spans their full range, so its edges rest on single values. Depth is given in mapped reads passing the MAPQ filter, the same unit as *n* in Table 2. Dashed lines mark the 0.6 and 0.8 thresholds of Mittnik et al. [10], a sample giving a value below the lower line may originate from a male and one above the upper line from a female individual.

Applying the thresholds of Mittnik et al. [10], misassignment was confined to the shallowest levels: 9 of 11,200 replicates were assigned to the wrong sex, all of them between 1,000 and 1,700 mapped reads (5 in the marten, 4 in the fox). From 2,000 mapped reads upwards, not one replicate of either sex or species was misassigned. The usual failure at low depth was instead non-assignment, where the confidence interval straddles the 0.6–0.8 corridor; its rate falls steadily with depth, from about a third at 1,000 reads to below 10% above 5,000, below 5% above 7,500 and below 1% above 10,000 in the marten and 15,000 in the fox. Both sexes behave alike. We report rates rather than the depth at which no replicate is left unassigned, because that depth is not a property of the method: more replicates surface rarer events and push it upward. With 200 replicates per level, a rate near 1% is resolved.

Simulated females did not centre on 1.00. At the deepest level their median *R*_*x*_ was 0.966 in the marten and 0.958 in the fox, and the male medians, 0.482 and 0.479, were almost exactly half of these, indicating an X-specific deficit of roughly 4% common to both sexes. The deficit is created by the MAPQ filter. Mapping a complete simulated female pool without any mapping-quality threshold gives an X-to-autosome density ratio of 1.000 in both species, the value the synthetic genomes prescribe; raising the threshold to MAPQ ≥ 30 retains 93.7% of the X-mapping reads against 97.3% of the autosomal ones in the marten, and 92.0% against 96.4% in the fox, bringing the ratio to 0.963 and 0.955, close to the replicate medians. The effect scales with the threshold rather than being peculiar to this one: at MAPQ ≥ 10 and ≥ 20 the marten ratio is 0.981 and 0.967. The X therefore carries a slightly smaller share of uniquely mappable sequence than the autosomes, and since the same filter is applied to the empirical samples, the simulated centre rather than 1.00 is the value against which they should be read. The medians are stable across depth, so this is an offset and not a depth-dependent bias.

From the simulation results, we adopt a two-step rule for empirical samples. (a) Depth: a sample is assigned only if it carries at least 2,000 reads mapping to the host chromosomes at MAPQ ≥ 30, the floor above which no simulated replicate of either species or sex was misassigned; below it a sample is reported as not assignable, whatever its confidence interval. (b) Interval: a sample passing step (a) is classified by the criterion of Mittnik et al. [10], as male if the upper bound of the 95% confidence interval is below 0.60, as female if the lower bound exceeds 0.80, and as unassigned otherwise.

### 3.3 *R*_*x*_ on the seven faecal metagenomes

Table 2 reports the empirical *R*_*x*_ values. The number of host-assigned reads after competitive mapping and MAPQ filtering ranged over almost three orders of magnitude, from 596 (sample 3) to 348,750 (sample 4). Four samples were classified as XX (1, 2, 4 and 5) and two as XY (6 and 7). In every assigned sample the Spearman correlation between chromosome length and mapped read count was high (0.88–1.00), indicating that read counts scaled with chromosome length as expected. Samples 6 and 7, assigned to males here, were not called male by *R*_*y*_ (Table 1).

**Table 2.** Sex estimates for the depositing individuals. *R*_*x*_ is the mean of the length-normalised ratios of reads mapping to the X chromosome and to the autosomes, *n* is the number of reads mapping to the chromosomes of the host species after competitive mapping (MAPQ ≥ 30), *ρ* is the Spearman rank correlation between chromosome length and read count. Assignment follows the two-step rule of Section 3.2: (a) *n* ≥ 2, 000; (b) following Mittnik et al. [10], male if the upper bound of the 95% confidence interval is *<* 0.60 and female if the lower bound is *>* 0.80. Sample 3 fails at step (a) although its interval would satisfy step (b).

| Species | Sample | *n* | *R<sub>x</sub>* | 95% CI | *ρ* | Assignment |
|---|---|---|---|---|---|---|
| *M. foina* | 1 | 8,479 | 1.06 | 1.00–1.12 | 0.97 | XX |
|  | 3 | 596 | 0.94 | 0.83–1.06 | 0.80 | n.a. |
|  | 4 | 348,750 | 0.96 | 0.94–0.98 | 1.00 | XX |
|  | 6 | 14,497 | 0.51 | 0.50–0.53 | 0.95 | XY |
|  | 7 | 4,191 | 0.39 | 0.37–0.41 | 0.91 | XY |
| *V. vulpes* | 2 | 50,746 | 0.98 | 0.97–1.00 | 0.98 | XX |
|  | 5 | 3,039 | 0.90 | 0.84–0.96 | 0.88 | XX |

Sample 3 was left unassigned. Its point estimate and interval (*R*_*x*_ = 0.94, 95% CI 0.83–1.06) would nominally satisfy the female criterion, but with 596 host-assigned reads it fails step (a) of the rule and is not assessed at step (b); its Spearman correlation was also the lowest in the set (0.80). All other samples pass step (a). Samples 5 and 7 (3,039 and 4,191 reads) fall in the range where some simulated replicates were left unassigned but none was misassigned, and both gave intervals clear of the ambiguous corridor, sample 7 by the widest margin (*R*_*x*_ = 0.39, 95% CI 0.37–0.41). Sample 1 (8,479 reads) lies above the depth at which the non-assignment rate falls below 5%, and samples 6, 2 and 4 above the depth at which it falls below 1%.

The value for sample 1 (*R*_*x*_ = 1.06, 95% CI 1.00–1.12) lies above the simulated female centre of 0.966, but well within the simulated spread at comparable depth: at 7,500 mapped reads the female marten replicates ranged from 0.77 to 1.13, and 8% of them reached 1.06 or more. That its confidence interval excludes the simulated centre reflects the narrowness of the interval rather than a departure needing a biological explanation.

## 4 Discussion

The *R*_*y*_ results in Table 1 illustrate a general problem rather than a defect of the original method. *R*_*y*_ was designed for a reference assembly in which the Y chromosome is represented adequately enough that the number of Y-mapping reads is proportional to Y content. In *M. foina*, the assembled Y is 5.6 Mb, a small target dominated by repetitive sequence. In *V. vulpes* there is none to count at all. The underlying reason is well understood, as mammalian Y chromosomes are built largely of ampliconic and palindromic repeats that defeat short-read assembly, which is why the first complete human Y sequence required long-read technologies and appeared only recently [17–20].

Comparing the *R*_*y*_ estimates against the *R*_*x*_ assignments of Table 2, the pattern is systematic rather than merely imprecise: the two samples that *R*_*x*_ places furthest into the male range, 6 and 7 (*R*_*x*_ = 0.51 and 0.39), are the two that *R*_*y*_ declines to call male. The reason is the numerator. The number of Y-mapping reads is 0 to 25 across the five samples, and the two *R*_*x*_-male samples contribute 18 and 2 reads. At these counts, the normal approximation on which the interval rests breaks down. Sample 3 yields *R*_*y*_ = 0 with a standard error of 0, and sample 7 yields an interval whose lower bound is negative, neither of which is a meaningful statement about a ratio. Sample 4 shows the same problem from the other side. It carries 350,796 host-assigned reads, more than any other sample, yet only 25 map to the Y, and this near-absence produces its confidently female *R*_*y*_. A statistic that misses males is worse than one that declines to answer, because the error is silent. We have no independent sexing of these animals and therefore report the discordance rather than resolving it, but its direction is what the assembly predicts: a 5.6 Mb, repeat-rich Y offers few positions at which a read can be placed uniquely at MAPQ ≥ 30, so the reads a true male does contribute are the ones most likely to be filtered out. For readers facing a similar decision, the lesson is that a Y-based ratio inherits every weakness of the Y assembly it is computed against, and that in taxa whose references lack a well-assembled Y the X-to-autosome ratio is the safer statistic – the same conclusion reached for ancient elephant remains, where *R*_*x*_-type approaches were adopted for comparable reasons [11].

Non-invasive samples collected on footpaths are handled and surrounded by people, and human reads were present in every library. For most metagenomic questions this is a nuisance; for sex determination it is a direct source of bias, because human reads that map to conserved X-linked regions of a carnivore reference inflate precisely the numerator of the statistic. Competitive mapping against a combined host-plus-human reference is the standard mitigation [13], and we regard it as a prerequisite rather than a refinement: a read-ratio statistic computed from a single-reference alignment of non-invasive material cannot distinguish host sequence from the conserved human sequence that resembles it.

A second precaution costs nothing and is worth taking. Because *R*_*x*_ averages X-to-autosome ratios across autosomes, a single chromosome that attracts reads at a rate incompatible with its length – through a repeat, a mis-assembly, or similarity to an abundant non-target template – shifts the mean, and if the effect is large enough it can push a female toward the male range. Our use of a rank correlation rather than a linear regression makes the diagnostic robust to such a chromosome, but it does not remove it from the average. We therefore recommend inspecting the per-chromosome counts before accepting an *R*_*x*_ value. The implementation reports them alongside the estimate.

The simulations answer one specific question: if reads are drawn at random from a genome with a known sex-chromosome complement and sequenced with the error profile of our run, do the human-derived thresholds of 0.6 and 0.8 classify the sample correctly at the read depths we actually obtain? The answer is yes, with a margin, as misassignment occurred in 9 of 11,200 replicates, all of them below 2,000 mapped reads. Above that depth the only failure mode was refusal to classify. This is the desirable failure mode for a monitoring application, where a missing value is far less damaging than a wrong one. In simulation, a floor of 2,000 host-assigned reads removes misassignment entirely, and non-assignment falls below 5% above 7,500 reads and below 1% above 15,000 reads in both species and sexes.

The simulations do not establish that the method is correct on real faecal DNA, and the gap between the two is worth stating plainly. Simulated reads are drawn from an error-free synthetic genome with uniform coverage expectation, whereas real faecal host DNA is fragmented, may be chemically damaged in the patterns characteristic of degraded templates [31], competes with an overwhelming microbial background, and comes from an individual that differs from the reference by structural variants and copy-number differences, a further source of reference bias [16]. Residual human reads that survive competitive mapping were not modelled at all. Each of these effects increases the variance of the per-autosome ratios, and greater variance widens the confidence interval. In the Mittnik framework, this pushes samples towards non-assignment rather than towards error, which is reassuring in direction but is an argument, not a measurement. A study in which scats of known-sex captive animals were processed through the same pipeline would convert this argument into evidence, and we regard such a validation as the natural next step.

Because VulVul3 contains no Y chromosome, our synthetic “male” fox genome consists of two autosome sets and a single X, with the Y simply absent rather than present but unassembled. For the *R*_*x*_ statistic, which does not use the Y at all, this is close to adequate: the X-to-autosome ratio of such a genome is the same as that of a true male. It is not identical in one respect, though, since in a real male a proportion of reads originate from the Y and are therefore not available to map elsewhere, whereas in our synthetic male those reads are drawn from the remaining chromosomes. The Y chromosome is a small part of the genome, so its effect is correspondingly small. However, this is a systematic rather than random difference, and it should be borne in mind when the fox simulations are used as a reference.

Four of the seven scats were assigned to female and two to male animals. The single unassigned sample failed because of too few host-assigned reads. We have deliberately not treated sample 3 as female even though its confidence interval nominally satisfies the criterion, because its 596 reads fall below the floor the calibration establishes. Its Spearman correlation, the weakest in the set at 0.80, points the same way. Reporting it as unassigned rather than as a weakly supported female is the conservative choice and is consistent with how we would wish such results to be used downstream.

None of the seven assignments has independent confirmation. No sex-linked PCR was run, no animal was observed, and the samples cannot be revisited. The assignments should therefore be read as the output of a calibrated but externally unvalidated procedure. Their internal consistency is encouraging – high Spearman correlations, agreement with the simulated centres, and clean separation between the two male samples (*R*_*x*_ = 0.39 and 0.51) and the four female samples (*R*_*x*_ = 0.90 to 1.06) – but internal consistency is not validation.

## 5 Conclusions

For low-coverage, human-contaminated faecal metagenomes of stone marten and red fox, the X-to-autosome read ratio *R*_*x*_ is a workable sex-determination statistic, while the Y-based *R*_*y*_ is not, for the simple reason that the available reference assemblies do not represent the Y adequately – 5.6 Mb of repeat-rich sequence in *M. foina* and nothing at all in *V. vulpes*. Competitive mapping must remove human contamination before computing the ratio, and per-chromosome read counts should be inspected. Under simulation with the error profile of a real sequencing run, the human-derived thresholds of 0.6 and 0.8 transferred to both species, misassigning 9 of 11,200 replicates, all below 2,000 mapped reads. We therefore report samples below that depth as unassigned, and expect non-assignment in under 5% of samples above 7,500 reads and under 1% above 15,000. Applied to seven scats, the procedure assigned four females and two males and declined to assign the seventh. The floor is specific to these two species, their assemblies, and a MAPQ 30 filter, and has to be re-established for another combination, for which the calibration script accompanying the implementation is intended. External validation on samples from animals of known sex remains outstanding and is the obvious next step.

## Declarations

### Data availability

The raw short-read data are publicly available through BioProject PRJNA1520986 in the NCBI Sequence Read Archive (SRA).

### Code availability

The source code of karyoratio, including the script that builds the chromosome table from an NCBI sequence report and the calibration script, is available at https://github.com/solymosin/karyoratio under the MIT licence. The version used here is tagged v2.0.0.

### Ethics statement

No animals were handled, captured or disturbed. All material consisted of faeces deposited naturally on publicly accessible hiking trails.

### Author contributions

Á*MA*: Review & Editing; *CM* : Formal analysis, Review & Editing, Writing – original draft; *IC* : Funding acquisition, Review & Editing; *GH* : Formal analysis, Review & Editing; *GM* : Investigation, Review & Editing; *NS* : Conceptualization, Formal analysis, Methodology, Review & Editing, Software, Supervision, Validation, Visualization, Writing – original draft; *VV* : Formal analysis, Review & Editing, Writing – original draft. All authors read and approved the final manuscript.

